# A Method for Optimizing Imaging Parameters to Record Neuronal and Cellular Activity at Depth with Bioluminescence

**DOI:** 10.1101/2023.10.29.564606

**Authors:** Alexander D. Silvagnoli, Kaylee A. Taylor, Ashley N. Slaviero, Eric D. Petersen

## Abstract

Optical imaging of activity has provided valuable insight into brain function and accelerated the field of neuroscience in recent years. Genetically encoded fluorescent activity sensors of calcium, neurotransmitters and voltage have been tools of choice for optical recording of neuronal activity. However, photon scattering and absorbance limits fluorescence imaging to superficial regions for *in vivo* activity imaging. This limitation prevents recording of population level activity in lower brain regions of experimental animals without implanted hardware. Single and multiphoton methods find maximal use in the cortex and experience loss of signal at greater depths. Successful efforts have been made to increase the depth of fluorescence imaging using fiber photometry and gradient reflective index lenses. However, these methods are highly invasive, requiring an implant within the brain. Bioluminescence imaging offers a promising alternative to achieve activity imaging in deeper brain regions without hardware implanted within the brain. Bioluminescent reporters can be genetically encoded and produce photons without external excitation. The use of enzymatic photon production also enables prolonged imaging sessions without the risk of photobleaching or phototoxicity. These characteristics render bioluminescence suitable to non-invasive imaging of deep neuronal populations. To facilitate the adoption of bioluminescent activity imaging, we sought to develop a low cost, simple *in vitro* method to optimize imaging parameters for determining optimal exposure times and optical hardware configurations to determine what frame rates can be captured with an individual lab’s imaging hardware with sufficient signal-to-noise ratios without the use of animals prior to starting an *in vivo* experiment. To achieve this, we developed an assay for modeling *in vivo* optical conditions with a brain tissue phantom paired with engineered cells that produce bioluminescence. We then used this assay to limit-test the detection depth vs maximum frame rate for bioluminescence imaging at experimentally relevant tissue depths using off the shelf imaging hardware. With this method, we demonstrate an effective means for increasing the utility of bioluminescent tools and lowering the barrier to adoption of bioluminescence activity imaging with bioluminescent sensors.

## Introduction

Both single and multiphoton fluorescence imaging suffer difficulties when acquiring signal from deep within the brain. Single photon imaging is typically restricted to superficial layers of cortex, with a recommended max signal resolution depth of <200 µm^1,2^. Beyond this depth, true signal becomes difficult to resolve from background values due to scattering, autofluorescence and photon energy loss. Multiphoton microscopes enable imaging at greater depth while achieving single cell resolution. Two and three photon imaging methods can achieve depths of 1,200 µm and 1700 µm respectively while providing single cell resolution^3-5^. Two-photon microscopy is well-suited for imaging the cortex but experiences significant scattering effects as a function of depth^6,7^. While 3-photon microscopy has the capability to image deeper than the cortex, it’s worth noting that the associated equipment costs may be challenging for many labs to accommodate. For these reasons, visualization of sub-cortical neuronal population dynamics continues to prove challenging or inaccessible for many labs.

Achieving optical recording in deeper brain regions has been achieved with fluorescence using techniques such as fiber photometry without single cell resolution. By exciting the genetically fluorescent sensor through a light fiber, activity can be recorded from a group of neurons near the tip of the fiber as a bulk population signal. This increases accessibility to deeper regions, however, comes at the cost of significant disturbance to brain tissue as well as limited population sampling^8,9^. The population size available for fiber photometry is limited to cells near the tip of the fiber. Pisanello et al. (2018) demonstrated that the effective volume of recording for a standard Ø400 µm, 0.50 NA optical fiber was a 10^6^ µm^3^ extending 200 µm in front of the fiber^10^. A fiber of similar dimensions to the provided example would be expected to displace a volume of ∼0.25– 0.5 mm^3^ when inserted between 2 – 4 mm^3^ deep respectively^11^. Also worth consideration are the effects of an implant within the tissue of a freely moving animal. The fiber employed for this type of imaging is typically stiffer than the surrounding tissue with few notable exceptions^12,13^. This resistance of the fiber to movement of the surrounding tissue can introduce disruptive forces to local vasculature, affecting both the blood brain barrier and parenchyma^12,14^. In freely behaving animals, this can introduce damage that cannot be accounted for until the end of the experimental period, especially for animals used for repeated recordings over time.

Gradient Refractive Index lenses, or GRIN lenses have gained popularity for activity imaging for their advantages over fiberoptic implants in providing single cell resolution. These lenses focus light by changing the refraction of light within the lens, providing a focused beam of photons on the subject^15,16^. However, they are not without limitations. The implantation of a lens is no less invasive than optical fiber. Recording from a larger population of neurons can require a lens of 0.5 – 2mm in diameter, displacing a tissue volume of ∼1.6 - 25 mm^3^ with the lens alone^17^. However, the total amount of tissue displaced is 2.8x greater than the volume of the lens. This is because the wall thickness for the glass tube accompanying the lens displaces yet more tissue. For example, the guide cannula for a 0.5 mm GRIN lens can be 0.84 mm in diameter^18^. As is a common drawback to most brain implants, implanted fibers and lenses can induce glial scarring, a condition where reactive astrocytes can encapsulate the implant, ultimately decreasing signal quality^19,20^.

Bioluminescence imaging offers an opportunity for deep brain activity data collection without the use of implants^15,21^. Both terrestrial and marine versions of luciferase enzymes are used in a wide range of life science fields including neurobiology and oncology^16,22,23^. Bioluminescent light is produced by enzymatic oxidation of a substrate, referred to as a luciferin^24^. By generating light within the tissue, it is possible to collect signal from deep structures without an implant, generally at the cost of lower spatial and temporal resolution compared to fluorescence imaging. Additionally, the production of light via consumption of substrates eliminates the possibility of photobleaching and has not been shown to have potential for phototoxicity. Thus, long imaging periods over repeated sessions can be leveraged to track long-term activity, a long-standing goal of neuroscience. Bioluminescent indicators can also be selectively expressed utilizing selective delivery methods like cell type specific or neuron subtype specific promoters and selective viral capsids. The specificity of expression coupled with in situ light generation simplifies the imaging requirements for capturing high signal to noise data, obviating the need for sophisticated optics due to the low hardware requirements for bioluminescence imaging^25^. Significant developments in the field of bioluminescence imaging have enabled signal detection from deep structures such as the hippocampus,^15,21^ basal ganglia^16,22,23^, hypothalamus^26^, and spinal cord^27,28^. Bioluminescence has also been imaged from implanted cells in freely behaving marmosets without surgical implants or attached hardware at a depth of 4.8mm^29^ and enabled bioluminescence based voltage and calcium imaging in freely behaving mice without implants or any attached hardware^30,31^.

Recently bioluminescent sensors have been developed that are capable of reporting calcium, voltage and neurotransmitter release, events that are crucial to neuronal communication and information processing. Genetically encoded bioluminescent indicators are favorable for deep tissue imaging due to the lack of endogenous light emission in mammalian tissue. This means that a relatively low photon output should be able to provide high SNR for transient calcium events. For example, Oh et al. (2019) utilized a red shifted bioluminescent calcium indicator (CamBI) to image activity of whole liver, imaging in the intact animal^32^. Tian et al. (2022) was able to record calcium activity in the basolateral amygdala through the closed skull utilizing their red shifted bioluminescent calcium sensor BRIC with a modified luciferase pro substrate, achieving bioluminescent activity imaging at a depth of 5 mm which is far below the effective fluorescence imaging range without an implanted lens or fiber^33^. Lambert et al. (2023) recently developed a green bioluminescent calcium sensor, CaBLAM with unparalleled dynamic range that we expect to enable more robust responses to be optically recorded in deep brain regions^34^. We have also recently expanded genetically encoded bioluminescent sensors to include detection of neurotransmitters *in vivo*, developing BLING (BioLuminescent Indicator of Glutamate) that emits blue light and demonstrated its ability to detect seizure activity in the rat at a depth of 2 mm while imaging through the closed skull^35^.

The promise shown in previous applications of bioluminescence activity imaging is promising but a barrier to widespread adoption is changing imaging approaches to accommodate the relatively low photon output when imaging through tissue. The scattering and absorbing properties of brain tissue limit the ability of photons to travel from their point of origin unchanged^36^. Computational and empirical calculations of the mean free path (MFP) determine that for a photon of a given wavelength, the distance that photon can travel without obstruction is approximately 90 µm^37,38^. Interference accumulated as a function of distance results in scattering, limiting the ability to resolve individual sources. Deep imaging of tissue is therefore restricted to characterization of whole neural populations rather than individual cells due to scattering and absorption. To compensate for limited photon production and decreased photon escape with increasing target depth, exposure times are lengthened, sacrificing temporal resolution for increased SNR to achieve non-invasive deep brain activity imaging.

Here, we present a method for optimizing imaging parameters with bioluminescence using a tissue phantom adapted from Ntombela et al. (2020) that simulates the absorbing and scattering properties of brain tissue to enable adoption of bioluminescent sensors for activity imaging. We paired this with a biological light source and demonstrated the ability to optimize imaging parameters without necessitating the use of animals to determine the detection limits of imaging hardware. This approach can be used to determine what timescale can be recorded using various imaging setups a lab may have to facilitate *in vivo* experimental design. Our goal is that researchers will be able to use this approach to determine what types of neuronal events and timescales can be captured prior to initiating animal experiments using an inexpensive optical brain phantom. We translate *in vitro* imaging conditions to *in vivo* like conditions, enabling approximation and optimization of *in vivo* imaging without significant animal use. To validate this approach, we employ the same imaging parameters *in vivo*. From these results, we expect that it is possible for others to easily adopt this approach for imaging hardware optimization and determine if physiological timescales can be recorded that align with experimental goals prior to initiating *in vivo* experiments, saving time, resources and reducing animal use.

## 2. Materials and Methods

### 2.1 Optical Tissue Phantom

We chose an optical phantom recipe from a previous publication by Ntombela et al. that was designed with the intent of replicating the light absorbing properties of multiple kinds of soft tissue. Their phantom was tested with multiple component ratios and validated with laser absorbance^41^. We used the ratio of components best suited for replicating brain grey matter optical properties. 2 grams agar (BD Ref 214530) and 300 mg of aluminum oxide [Al_2_O_3_] (Sigma Aldrich, 199443) were weighed and mixed in an Erlenmeyer flask containing 100 mL de-ionized water and a magnetic stir bar. The mixture was heated while stirring until the temperature reached 94°C. Temperature was monitored continuously using ThermPro Food Thermometer (Model No. TP-02). Once reached, the mixture was moved to a separate stir plate at 55° C to prevent early firming. 300 µL India ink (Bombay, BOMB10OZS7BY) was added and stirred until completely mixed.

The mixture was then pipetted into 96 well plates for molding. Plates containing molten phantom were covered with parafilm and the original lid to prevent loss of moisture while firming. Each plate was placed at 4°C to rapidly solidify to prevent precipitation of the aluminum oxide and left for at least 24 hours. Agar based brain phantoms in the shape of pucks were later removed from the plates via hydraulic pressure forced from the bottom of the well. To accomplish this, a 30-gauge hypodermic needle was attached to a 3 mL syringe containing 1x phosphate buffered saline (PBS). The needle was inserted at the periphery of a phantom puck, carefully not puncturing the puck and PBS was injected into the bottom of the well until the puck was fully ejected. Using this technique, we were able to leverage the generated pressure to force the puck out of the well without damaging it.

The solidified phantom pucks with lengths of 2, 4, and 6 mm were cut using a spinal cord steel matrix (Alto SA-5110) and stainless-steel stainless single edge razor blades (VWR 55411-050). This instrument was utilized simply due to already having it and we expect many other means can be used to accomplish this task with sufficient precision. To control potential variability, pucks for a given ‘depth’ were weighed, and a running average was established. This average was later used to preemptively remove potential outliers. Pucks differing by more than 2 mg from the average for a given ‘depth’ were disposed of prior to imaging experiment.

### 2.2 Evaluating the Optical Tissue Phantom against Brain Tissue

Pucks of 2, 4 and 6 mm depths were cut and placed into black clear bottom 96 well plates (Corning Ref. 3603). Experiments were performed using replicates of 5 per given depth. Ink concentration was varied according to a 1:2 serial dilution for comparison of ink concentration vs depth.

Brain tissue was harvested from mice C57BL/6 wild type mice (age > P60) with sufficient tissue to run absorbance replicates in triplicate. Animals were euthanized using CO_2_ followed by cervical dislocation, the brain immediately removed and put in PBS chilled on ice. Mice were deliberately not perfused. Brain tissue samples were divided into three groups: unfixed, 24-hour fixation, and 48-hour fixation. Fixed brains were placed in 4% para-formaldehyde in PBS at 4°C. Unfixed brains were extracted and used same day. Brain tissue samples were sectioned into lengths of 2, 4, and 6mm. Brain samples were placed into wells in replicates of three alongside the phantom pucks. Wells containing PBS were used as a blank control.

Absorbance spectra were generated using a multipoint 3×3 well scan of each well from 450 to 700 nm with a plate reader (Tecan Spark). Preliminary experiments determined that there was a significant difference in absorption by the phantom pucks when the absorbance was read at room temperature (approx. 27°C) and 37°C.

### 2.3 Bioluminescence imaging through the Phantom

#### 2.3.1 Acute Lipofection with SSLuc

Hek-293 cells were in a 24-well plate, grown to 70% confluency. Each well was then treated with Lipofectamine 2000 (Thermofisher Cat No. 11668-019) [2 µL/well] and SSLuc plasmid [0.8 µg/well]. Media was changed after 4 hours and cells incubated for 48 hours before testing.

#### 2.3.2 IVIS Imaging: Bioluminescent Cells

Acutely transfected HEK293 cells expressing SSLuc were suspended and concentrated in FluroBrite DMEM to 6×10^6^ cells/mL (Ref A18967-01). 35 µL of cells (∼200,000) were added to wells spaced two rows and two columns apart. 5 µL of 2 mM hCTZ was added to wells containing cells just prior to being placed in an IVIS Lumina. Pucks produced as described previously were placed over wells containing cells in the IVIS machine and imaged at 100, 250, 500, 1000, and 2000 ms exposure times. The binning was set to medium, and the F-stop was set at 1. The height of the camera was set to 1.5 cm. Temperature for the IVIS was set at 37°C.

#### 2.3.3 Stable cell line generation

Flp-In-293 cells (ThermoFisher Scientific Ref R75007) were transfected according to standard lipofection procedure (Lipofectamine 2000 ThermoFisher Scientific) with a plasmid containing the intact bioluminescent luciferase SSLuc that was used to create the calcium sensor CaBLAM and a Hygromycin B resistance gene^34^. Cell media was changed 6 hours after transfection and cells were split the next morning and seeded at ∼25% confluency on a new plate. Three days post transfection, cells were treated with Hygromycin B (Invitrogen Ref. 10687010) [100 µg/mL] with DMEM, 10% FBS, 1x Pen/Strep. Media was changed and fresh Hygromycin B was added every three days until cell colonies had formed. Hygromycin resistant colonies were pooled and expanded. Single cells were then selected by performing a dilution series on the polyclonal pool and seeding approximately one cell per well on a 96-well plate. Cultures derived from monoclonal colonies were evaluated in replicates of 5 in 96 well plates for consistency of expression and brightness in response to h-Coelenterazine (h-CTZ) [250 µM] (NanoLight, Ref #301) administration, read with a Tecan Spark plate reader.

#### 2.3.4 Bioluminescence Microscope Set-Up

We assembled off the shelf components likely to be standard equipment for many labs. We mounted an Andor EMCCD camera, xIon 888 onto a dissection microscope (Leica No. MZ10 F) with 1.0x objective and a tube lens adaptor (Leica No. 10447367). Our set up was surrounded by a home-made dark box constructed of 1/2-inch PVC tubing found at most hardware stores, covered in black drop cloth (Thor Labs Ref. BK5). To ensure the dark box was not permissive of light, we tested the inside of the box using a light power meter (Thor Labs Ref. S121C). Recorded values were found to not be significantly different from background noise of the light power meter when covered and room lights off. This was interpreted to mean that the dark box surrounding the microscope was light tight.

#### 2.3.5 Bioluminescence Microscope Imaging: Bioluminescent cells

Confluent Flp-In-SSLuc cells were suspended and concentrated as described above. hCTZ was also prepared as described above. Phantom pucks of 2, 4, and 6 mm were cut and placed in PBS to prevent dehydration. Prior to recording, 100 frames of dark background were recorded for later analysis. Wells were centered in the field of view and an image was taken to use as a reference for ROI placement. Each well was prepared and recorded individually. Recorded wells were spaced two wells apart vertically and horizontally. Stocks of cells and hCTZ were generated and kept at 37 °C in a warm water bath for the duration of testing. 30 µL of cells and 30 µL of hCTZ [0.5 mM] were mixed in microcentrifuge tubes and pipetted up and down multiple times to ensure even mixing. 40 µL of this mixture was placed into a well and covered immediately with a phantom puck. The drape covering the light proof box over the microscope was secured over the dark box. Each sample was recorded for 200 frames at 95, 100, 150, 250, 500, 1000, and 2000 ms exposures.

### 2.4 In Vivo Data Collection

#### 2.4.1 Viral Transduction

C57BL/6 mice between 4 – 8 months were injected unilaterally using AAV-*hSyn*-ssLuc at stereotaxic coordinates (-2.5, -1) relative to bregma. Prior to surgery mice were anesthetized using isoflurane at 2.0%. A small burr hole was created above the coordinates using a dental drill. Using a World Precision Instruments syringe with 33-gauge needle (NanoFil Ref. NF33BV), mice were injected with 1 µL of virus [3×10^13^ vg/mL] at 2 or 4 mm of depth relative to the top of the cortex at a rate of 0.5 µL/min. Following completion of the injection, the needle was allowed to sit at depth for 10 minutes. A small dab of dental cement was used to seal the burr hole and the skin was closed with sutures. Mice were treated with triple-antibiotic ointment and lidocaine.

#### 2.4.2 Skull Thinning

After two weeks, the skull covering the injected hemisphere was thinned. Mice were anesthetized with isoflurane. A dental drill was used to gently shave away bone tissue from the injected side until only cortical bone remained^42^ and wound sutured closed. Mice were allowed to recover from surgery in a heated chamber at 37°C until grooming was observed. A 48-hour recovery period followed the procedure.

#### 2.4.3 Animal Imaging with IVIS

Thirty minutes prior to anesthesia, animals were individually injected IP with water soluble hCTZ (NanoLight Cat#3011). Each was anesthetized using a ketamine (10 mg/kg) and xylazine (2 mg/kg) cocktail and tested for toe and tail pinch response. The skin above the thinned skull was re-opened and pulled to either side using surgical clips. Emitted bioluminescence was recorded through the exposed, thinned skull using the same parameters as the plates with phantoms at 100, 250, 500, 1000, and 2000 ms. Animals were then allowed to recover in a heated recovery chamber until alert.

#### 2.4.4 Animal Imaging with EMCCD

Similar to the IVIS imaging, animals were injected with hCTZ (10 mg/kg) and then anesthetized with a ketamine/xylazine cocktail. After checking for toe and tail pinch response, the thinned skull above the injection site was exposed and surgical clips were used to pull the skin aside. Mice were placed on a heated pad and bioluminescence was recorded at the exposure times specified in (Fig 5C). A long exposure was taken at the end of the recording period to validate the presence of bioluminescence. Mice were perfused immediately at the end of their recording session. All procedures were approved by Central Michigan University IACUC.

## 3. Results

### 3.1 Absorption Spectra of Phantom vs. Brain Tissue

By measuring the absorbance of the brain phantoms, we determined the ink concentration reported to correspond to brain tissue in Ntombela et al. (2020) closely matches the absorbance curve we generated, compared to freshly harvested mouse brain tissue but not fixed brain tissue (Fig 1A)^41^. Interestingly, because our coverage of the visible light spectrum included several points between those tested in Ntombela et al. (2020), we were able to observe a bump in the fresh tissue absorbance from 500 to 600 nm (Fig 1B). These results agree with the absorbance of hemoglobin in fresh tissue^43^. This was done to simulate *in vivo* optical conditions as closely as possible. Examination of the curves for fresh tissue and the 0.3% India Ink phantom shows a close overlap throughout most of the visible light spectrum. Analysis using 2-way ANOVA showed no statistically significant difference between the phantom and fresh tissue at the wavelengths of interest except for 470 nm (p < 0.01) (Fig 1C).

**Figure 1.**
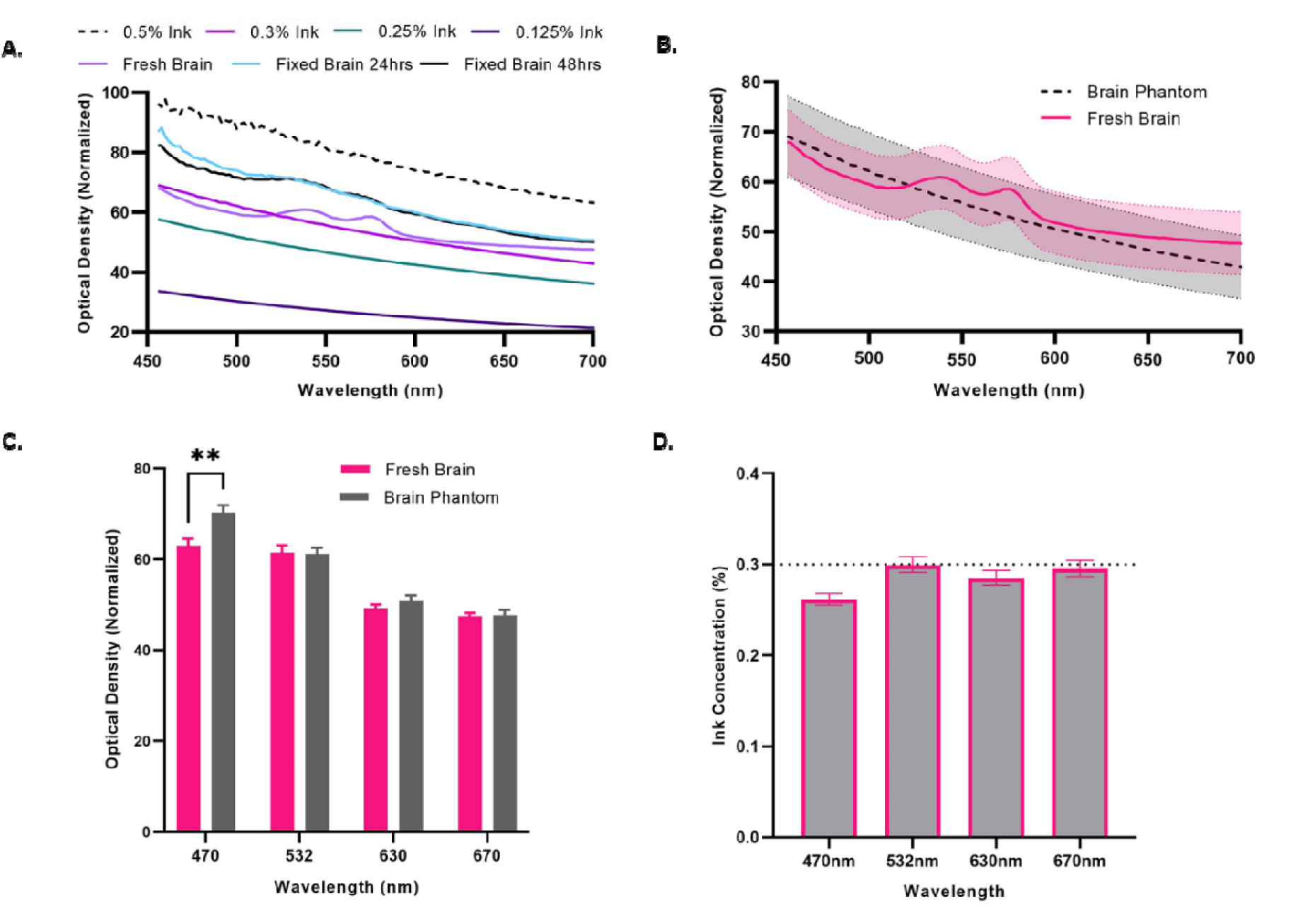
Evaluation of the Phantom as a Proxy for Brain Tissue. **A**. Plot of the absorbance of phantom pucks against wavelength. **B**. Absorbance plotted for the brain phantom vs. fresh tissue illustrates the overlap between fresh brain tissue and the brain phantom at 0.3% India Ink concentration. **C**. Comparing the absorbance values of fresh brain tissue vs. the phantom at 470, 532, 630, and 670 nm. No significant differences shown for each wavelength as determined with 2-way ANOVA except for 470 nm (p < 0.01). **D**. Interpolated ink concentrations of brain tissue based on the absorbance curves and regression modelling for the wavelength of interest. The dotted line indicates the ink concentration 0.3% reproduced from by Ntombela et al. (2020) to match the absorbance of grey matter.

To further validate that our findings were consistent with expectations, we back-calculated hypothetical ink concentrations based on the observed optical density (OD) of fresh tissue. Using non-linear regression modeling with Prism 9, the decay of optical density in the phantom as a function of increased wavelength was generated for the phantom containing 0.3% ink (R^2^ > 0.95). OD values of brain tissue at critical wavelengths were then interpolated along the curve to determine the approximate “ink concentration” required to match that of fresh brain tissue, of which 470nm would require significantly lower ink, expected to be 0.265 (Fig 1D). Here we found that it was possible to closely model the absorbance of light by brain tissue using the phantom alone. We believe this is significant due to the extreme simplicity of the phantom itself relative to the natural complexity of optimizing imaging experiments with animal models. This was encouraging as our aim was to reduce the overall time and financial cost of optically modeling the brain.

We combined our phantom with an opaque bottom black 96-well plate (Corning Ref. 3650) to simulate light generation from within the brain as closely as possible. Phantom pucks were placed on top of small volumes of media containing bioluminescent cells treated with hCTZ. Wells were tested in duplicates of 2, 4, and 6 mm puck depths with additional wells for blank (i.e. non-treated) controls (Fig 2a). Our data showed that it was possible to collect bioluminescent light through the tissue phantom at significant levels above background. However, 2-way ANOVA analysis of wells containing transfected HEK293 SSLuc cells treated with hCTZ did not show statistical significance against background until after 500 ms of exposure (Fig 2B). It is also interesting to note that these experiments also revealed that quantification of emitted photons was not possible at or below 250 ms with the conditions used in our experiment (Fig 2B).

**Fig 2.**
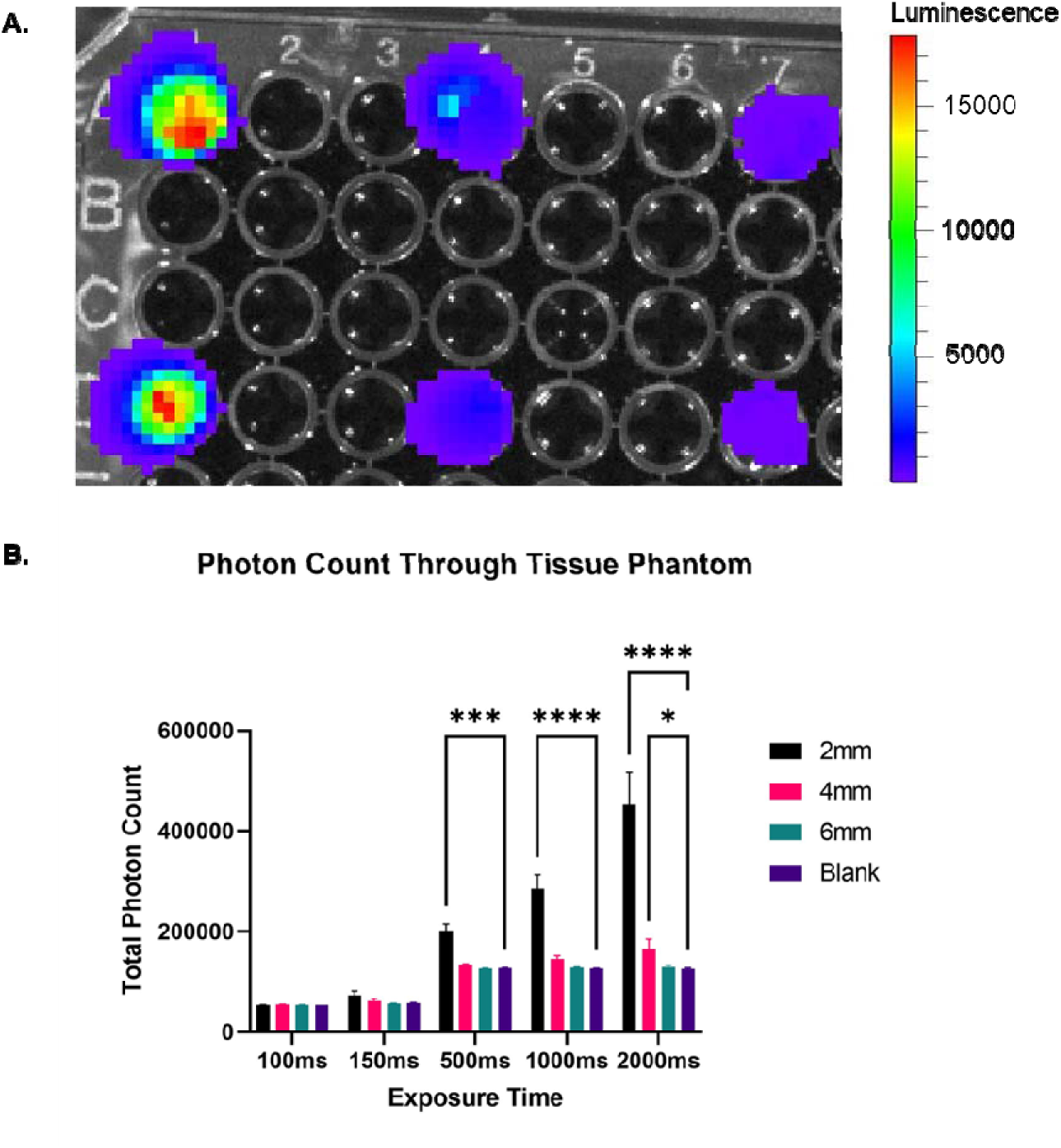
Testing the brain phantoms with an IVIS. A) A representative image of bioluminescence from our engineered cells recorded through phantoms of varying thicknesses. From left to right, the wells are occupied by 2, 4, and 6 mm phantom pucks. B) Measured values of photon count through the tissue phantom. Photons were generated from HEK293 cells expressing a green-shifted bioluminescent constructed emitting photons at the 535 nm wavelength.

### 3.2 Evaluation of IVIS Imaging System

We explored a wide array of imaging technologies. Options included custom microscopes, commercial options, and even lens-less imaging. A secondary goal of this project was the reduction of cost for imaging bioluminescence. We chose to test readily available and ubiquitously used imaging equipment to create our imaging setup Equipping a standard dissection microscope with an EMCCD camera and tube lens adaptor and building a light tight imaging chamber. Recording with this set-up followed a similar format to the IVIS experiments. Fig 3A. shows the workflow for this experiment. Fig 3B shows a representative image for a 4 mm phantom at 500 ms exposure. We found that using our set-up, it was possible to detect signal above background beginning at 95 ms exposure times (Fig 3C). With our equipment, this translated to a recording speed of 10Hz. The level of brightness above background was significantly greater for all exposures for the 2 mm phantom depth. As expected, 4 and 6 mm phantoms were much less permissive of light. Here we see that it is possible to detect relevant signal above background.

**Figure 3.**
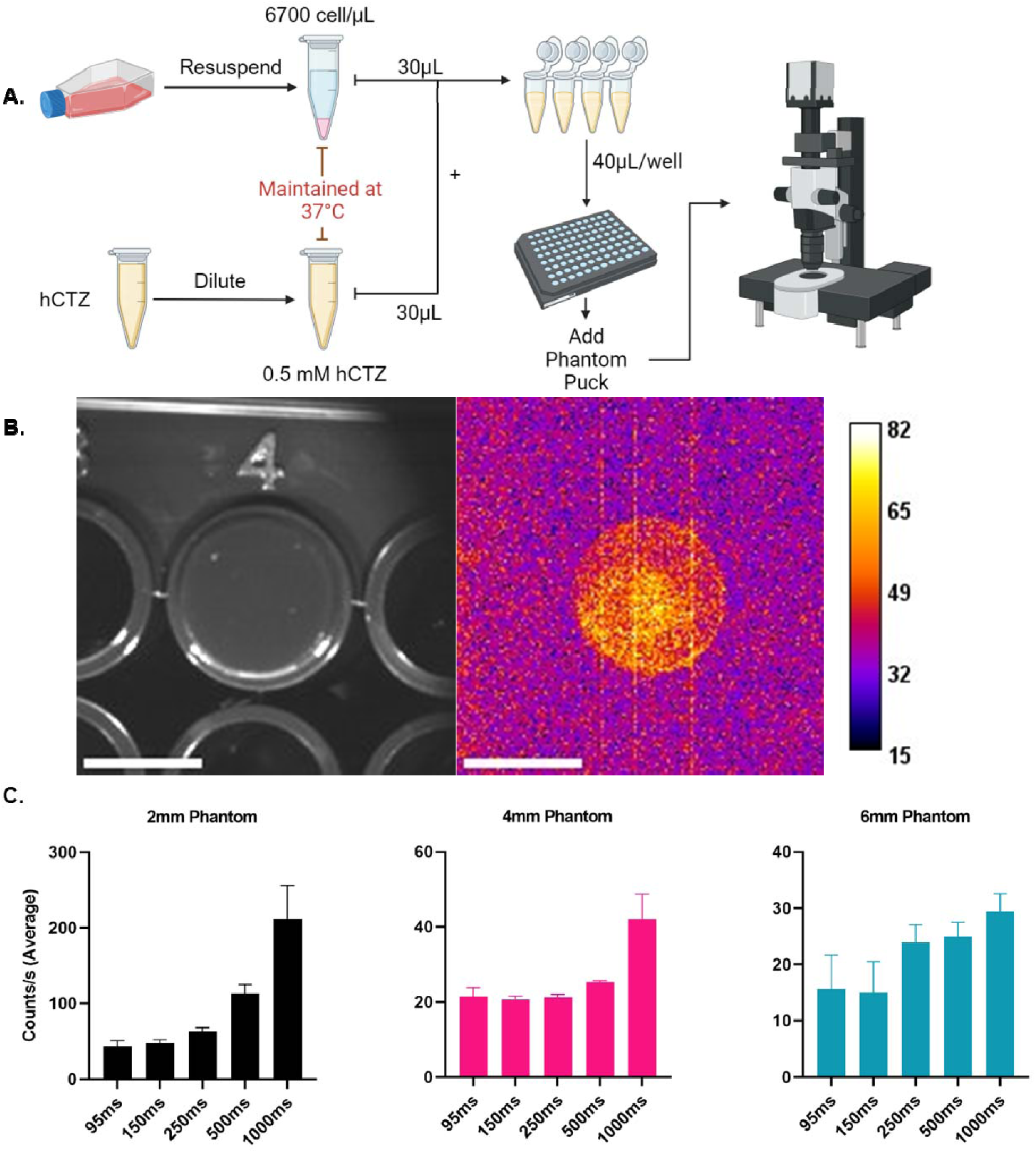
EMCCD Recording of Bioluminescence Through Tissue Phantom. **A**. A diagram of the workflow illustrating the method of bringing the cells from culture to testing **B**. A representative image showing the bioluminescence intensity (right) and brightfield (left) of a 4mm phantom puck in the 96-well black plate. **C**. Measured total photon emission was detected with an Andor EMCCD iXon camera at each exposure time. Each condition tested in replicates of 3. Counts calculated with averaged blank background subtracted from each recording.

### 3.3 Evaluation of EMCCD Imaging Set Up

To validate our findings from our *in vitro* model, we transduced mice at 4 mm with AAV9-*hSyn*-SSLuc. Total counts from the selected ROI with the IVIS were generally higher compared to the EMCCD setup when testing longer exposure times. However, the IVIS was not able to record significant signal above background at exposures shorter than 250 ms for either phantom or animals. These results closely match the phantom recordings from the SSLuc stable cell line.

**Figure 4.**
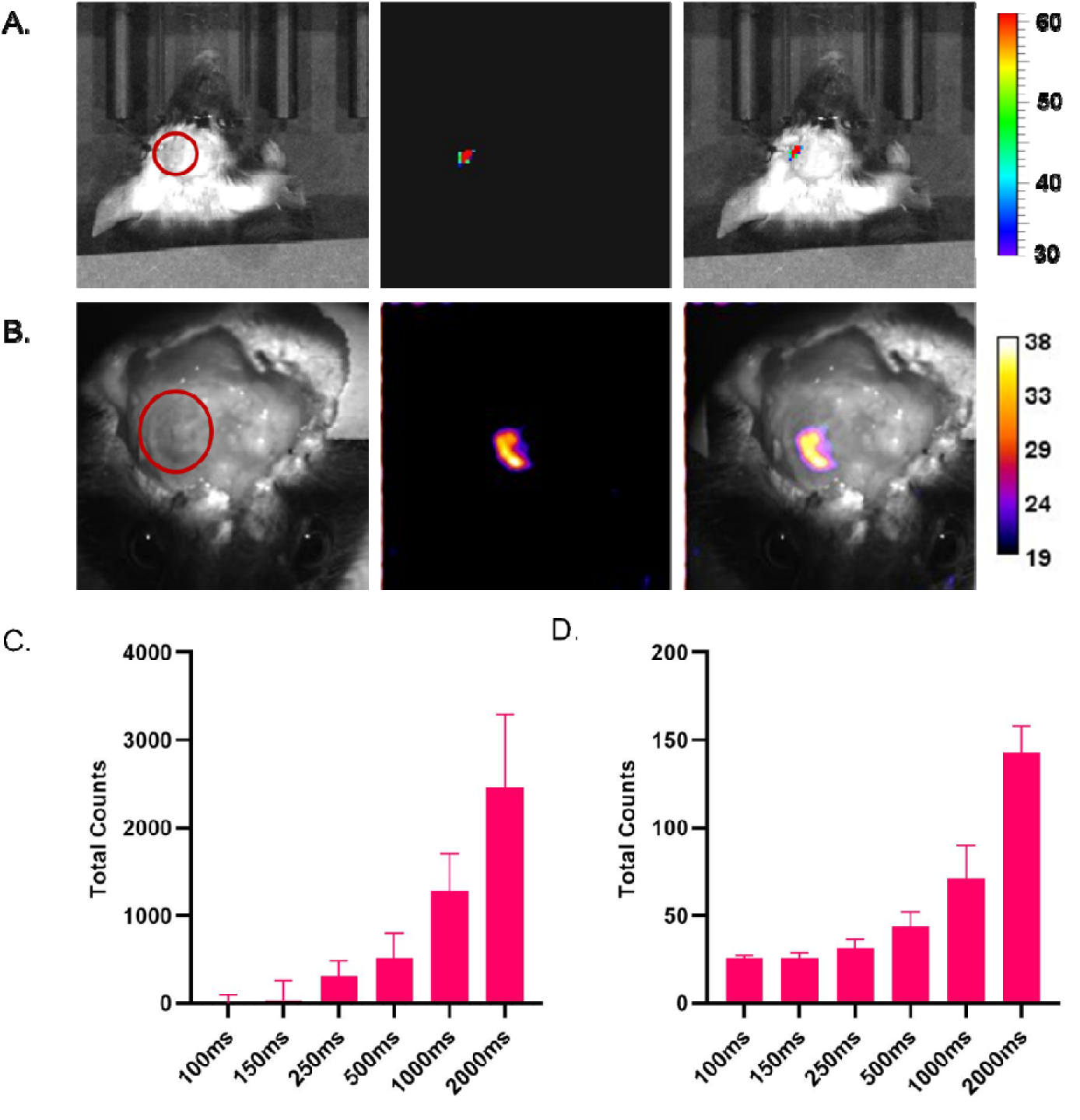
Mice transduced with SSLuc at 4 mm depth imaged with IVIS and EMCCD. **A**. and **B**. Red circles indicate where skull was thinned. **A**. Bioluminescence collected on the IVIS imaging system at 1000 ms exposure. **B**. Bioluminescence collected through thinned skull with the EMCCD camera at 95 ms exposure time. **C**. Total Counts of pixels recorded on the IVIS. **D**. The same measure collected with the EMCCD camera. Graph shows total counts averaged across animals (n = 3). Error bars indicate SEM.

## 4. Analysis

### 4.1 Absorption Spectra of Phantom vs. Brain Tissue

Absorbance spectra collected using the Tecan Spark was evaluated using GraphPad Prism Software. Replicates were checked for percent covariation to evaluate the quality of the collected data. 2-way ANOVA was used to compare optical density (OD) values at 470, 530, 630, and 670 nm.

### 4.2 Estimation of Analogous Ink Concentration

Absorbance data for 470, 530, and 670 nm wavelengths were fitted to linear regression plots. Area scans were performed using multipoint scan at frequencies 430, 560, 630 and 670 nm. Area scans were averaged for within well values and normalized to wells containing 1x PBS only. Replicates were analyzed for %CV for quality control. All wells had minimal variability and were included in analysis.

### 4.3 IVIS Image Analysis

Initial experiments were done with the IVIS due to its widespread use for imaging bioluminescence. Bioluminescence was measured from ROI’s matching the size of the well and expressed as total counts from the included Living Image software. Replicates (n = 2) were averaged for a given phantom depth for each exposure time. Values were analyzed with 2-way ANOVA and compared to blank wells (Fig 2B).

### 4.4 EMCCD Image Analysis

Background was recorded for each exposure time prior to recording bioluminescence for 100 frames (binning 4×4, Photon Counting preset). Image processing was performed using Fiji. Recorded frames were scanned for visible aberrations and each frame with a significant artifact such as a bright line across the frame was deleted from the stack. A gaussian blur was applied to each frame of the cleaned stack and then the stack averaged into a single image. Averaged dark frames were subtracted from experimental recording Z-stacks for a given exposure time. For example, the background averaged dark image was subtracted from each frame of a 95 ms exposure recording. Files were then filtered for outliers and then z-projected according to the stack average. The well in each image containing a sample was measured with an ROI corresponding to the size of the well. For *in vivo* experiments, ROIs were placed over the entire area of thinned skull. Replicate values for each depth were averaged across their exposure times. Representative images were pseudo-colored with a look up table and adjusted in brightness. Calibration bars were generated with Fiji and attached to representative images.

## 5. Discussion

In summary, we have developed a method for optimizing imaging conditions for recording bioluminescent light at a fast frame rate and at significant depth for labs to optimize hardware and imaging parameters prior to initiating *in vivo* experiments. We began by validating the brain tissue phantom for light absorbance compared to fixed and fresh brain tissue, testing different concentrations of absorbance material to validate the phantom’s use and build upon previous work. Each subsequent step of our experiments was tested in both our own microscopy set-up and the commercially available IVIS. We next tested a green light emitting luciferase expressed in HEK cells with this brain tissue phantom. These results were validated in mice transduced with the same construct. Bioluminescence imaged through thinned skulls have similar brightness to the *in vitro* stable SSLuc cells and phantom combination. We were able to record significant bioluminescent activity above background with high SNR.

In our study, we developed an *in vitro* method of modeling bioluminescent detection by simulating scattering and absorption using a previously characterized in brain tissue phantom. Part of the incentive to use this phantom was to reduce the number of animals required to determine ideal imaging conditions for a given bioluminescent construct. A key limitation in the adoption of new bioluminescent reporters is the transition from *in vitro* to *in vivo*. The simple fact is that modeling or creating expectations for the precise effects tissue will have on the reduction of collected photons for a newly designed construct is not realistic. Thus, functional testing and optimization throughput time can be increased with preliminary testing *in vitro* with simulated *in vivo* like conditions.

Previous studies have performed activity imaging with bioluminescent light emission recorded through the closed skull by red shifted luciferase/substrate variants. For example, work by Tian et al. (2022) generated a bioluminescent calcium sensor capable of reporting neuronal activity through closed skull. Calcium flux was detected and reported seizure activity in real time for areas of the hippocampus and responses to foot shock in the basolateral amygdala (5mm depth) at a frame rate of 1 Hz (1 s exposure). Oh et al. (2019) was able to report calcium fluctuations non-invasively in the liver using their bioluminescent orange calcium sensor. Images for these experiments were captured using the IVIS at 0.5 Hz (2 s exposure). These studies demonstrate that it is possible to detect relevant signals from bioluminescent reporters beneath dense tissue.

When running our experiments using a green-emitting bioluminescent construct (535 nm), we were able to effectively collect sufficient light above background. For imaging fast activity that occurs on time scales shorter than 250 ms, we found the EMCCD imaging setup shows better sensitivity, recording at 10Hz, 95 ms exposure plus ∼5 ms hardware/data transfer delay while in this case the IVIS successfully recorded at 4 Hz. One of our goals for this study was to effectively record at a frame rate close to or higher than 10 Hz. The discovery of this hardware limitation led us to seek out alternative equipment to incorporate as part of our new method. Utilizing an EMCCD camera and off-the-shelf microscopy components, we were able to overcome this recording limitation. In doing so we demonstrated that it is possible to record lower-than-red wavelength bioluminescent signal through an optically challenging medium at relatively fast frame rates.

To validate that our *in vitro* model could simulate the *in vivo* conditions with reasonable fidelity, we transduced mice at 4 mm with virus containing the SSLuc construct. In this case, we opted for skull thinning because it allows improved photon escape while still preserving the integrity of the BBB (blood brain barrier), the skull of transduced mice was very carefully thinned prior to imaging. We were able to preserve the integrity of BBB underneath, using the condition of the vasculature as a proxy for BBB health. Thinning the skull allowed detection signal at low exposure times with frame rates. This included a 95 ms exposure which resulted in a frame rate of 10 Hz including hardware delay. We therefore believe it is reasonable to expect that activity occurring on fast time scales, such as neurotransmitter release, could be detected with bioluminescence in real time when adopting custom built optical setups that we expect to allow detection of an order of magnitude more photons per exposure time^45,46^. This validation testing further solidified the expectation that this phantom method could be used to enhance a bioluminescent construct development pipeline and adoption of bioluminescence for routine activity imaging in labs while minimizing resources.

A limitation of our method is the simplicity of the phantom itself. As is visible in the absorbance curve of Fig 1., brain tissue is highly heterogenous with multiple tissue types and presence of hemoglobin contributing to its light absorbing properties varying with wavelength. The brain tissue phantom is consistent throughout and does not reflect this variability in tissue type. Future directions could explore different ways of layering the agar-based phantom we used as a part of our study. Additionally, the microscopy set up we used, could likely be improved upon with custom setups that are more permissive to photons. There were multiple lenses between the EMCCD sensor and the bioluminescent light source itself including the use of a zoom lens. This likely reduced the overall signal due to loss of photons as they traveled through the imaging path. Other groups have employed, custom built microscopes that we expect would be capable of achieving high frame rate bioluminescent light recordings^45-47^. A caveat to these designs is that they require calibration and a level of expertise in construction that many may not have the time or resources necessary to effectively utilize for initial adoption of bioluminescence imaging. For this reason, we chose to use off the shelf components including a pre-assembled dissection microscope for this study and expect to improve on our results going forward.

## References

1 Iwasaki, S. & Ikegaya, Y. In vivo one-photon confocal calcium imaging of neuronal activity from the mouse neocortex. Journal of Integrative Neuroscience 17, 671–678 (2018). 10.3233/JIN-180094

2 Yang, W. & Yuste, R. In vivo imaging of neural activity. Nat Methods 14, 349–359 (2017). 10.1038/nmeth.4230

3 Cheng, X. et al. Comparing the fundamental imaging depth limit of two-photon, three-photon, and non-degenerate two-photon microscopy. Opt Lett 45, 2934–2937 (2020). 10.1364/OL.392724

4 Ouzounov, D. G. et al. In vivo three-photon imaging of activity of GCaMP6-labeled neurons deep in intact mouse brain. Nature Methods 14, 388–390 (2017). 10.1038/nmeth.4183

5 Hontani, Y., Xia, F. & Xu, C. Multicolor three-photon fluorescence imaging with single-wavelength excitation deep in mouse brain. Science Advances 7, eabf3531 10.1126/sciadv.abf3531

6 Durr, N. J., Weisspfennig, C. T., Holfeld, B. A. & Ben-Yakar, A. Maximum imaging depth of two-photon autofluorescence microscopy in epithelial tissues. J Biomed Opt 16, 026008 (2011). 10.1117/1.3548646

7 Streich, L. et al. High-resolution structural and functional deep brain imaging using adaptive optics three-photon microscopy. Nat Methods 18, 1253–1258 (2021). 10.1038/s41592-021-01257-6

8 Nazempour, R., Zhang, Q., Fu, R. & Sheng, X. Biocompatible and Implantable Optical Fibers and Waveguides for Biomedicine. Materials (Basel) 11 (2018). 10.3390/ma11081283

9 Potter, K. A., Buck, A. C., Self, W. K. & Capadona, J. R. Stab injury and device implantation within the brain results in inversely multiphasic neuroinflammatory and neurodegenerative responses. Journal of Neural Engineering 9, 14 (2012). 10.1088/1741-2560/9/4/046020

10 Pisanello, M. et al. The Three-Dimensional Signal Collection Field for Fiber Photometry in Brain Tissue. Front Neurosci 13, 82 (2019). 10.3389/fnins.2019.00082

11 Taal, A. J., Lee, C., Choi, J., Hellenkamp, B. & Shepard, K. L. Toward implantable devices for angle-sensitive, lens-less, multifluorescent, single-photon lifetime imaging in the brain using Fabry–Perot and absorptive color filters. Light: Science & Applications 11, 24 (2022). 10.1038/s41377-022-00708-9

12 Sparta, D. R. et al. Construction of implantable optical fibers for long-term optogenetic manipulation of neural circuits. Nat Protoc 7, 12–23 (2011). 10.1038/nprot.2011.413

13 Cao, Y. et al. Flexible and stretchable polymer optical fibers for chronic brain and vagus nerve optogenetic stimulations in free-behaving animals. BMC Biol 19, 252 (2021). 10.1186/s12915-021-01187-x

14 Polikov, V. S., Tresco, P. A. & Reichert, W. M. Response of brain tissue to chronically implanted neural electrodes. Journal of Neuroscience Methods 148, 1–18 (2005). 10.1016/j.jneumeth.2005.08.015

15 Fricke, I. B. et al. In vivo bioluminescence imaging of neurogenesis - the role of the blood brain barrier in an experimental model of Parkinson’s disease. Eur J Neurosci 45, 975–986 (2017). 10.1111/ejn.13540

16 Aswendt, M. et al. Quantitative in vivo dual-color bioluminescence imaging in the mouse brain. Neurophotonics 6, 025006 (2019). 10.1117/1.NPh.6.2.025006

17 Wei, B. et al. Clear optically matched panoramic access channel technique (COMPACT) for large-volume deep brain imaging. Nature Methods 18, 959–964 (2021). 10.1038/s41592-021-01230-3

18 Bocarsly, M. E. et al. Minimally invasive microendoscopy system for in vivo functional imaging of deep nuclei in the mouse brain. Biomed Opt Express 6, 4546–4556 (2015). 10.1364/boe.6.004546

19 Kahan, A. et al. Light-guided sectioning for precise in situ localization and tissue interface analysis for brain-implanted optical fibers and GRIN lenses. Cell Rep 36, 109744 (2021). 10.1016/j.celrep.2021.109744

20 Xie, Y. et al. In vivo monitoring of glial scar proliferation on chronically implanted neural electrodes by fiber optical coherence tomography. Front Neuroeng 7, 34 (2014). 10.3389/fneng.2014.00034

21 Dunn-Meynell, A. A. et al. In vivo Bioluminescence Imaging Used to Monitor Disease Activity and Therapeutic Response in a Mouse Model of Tauopathy. Front Aging Neurosci 11, 252 (2019). 10.3389/fnagi.2019.00252

22 Kim, D. E. et al. Neural stem cell transplant survival in brains of mice: assessing the effect of immunity and ischemia by using real-time bioluminescent imaging. Radiology 241, 822–830 (2006). 10.1148/radiol.2413050466

23 Tung, J. K., Gutekunst, C. A. & Gross, R. E. Inhibitory luminopsins: genetically-encoded bioluminescent opsins for versatile, scalable, and hardware-independent optogenetic inhibition. Sci Rep 5, 14366 (2015). 10.1038/srep14366

24 Shimomura, O. Bioluminescence. (WORLD SCIENTIFIC, 2006).

25 Celinskis, D. et al. Miniaturized Devices for Bioluminescence Imaging in Freely Behaving Animals. Annu Int Conf IEEE Eng Med Biol Soc 2020, 4385–4389 (2020). 10.1109/embc44109.2020.9175375

26 Cordonier, E. L. et al. Luciferase Reporter Mice for In Vivo Monitoring and Ex Vivo Assessment of Hypothalamic Signaling of Socs3 Expression. J Endocr Soc 3, 1246–1260 (2019). 10.1210/js.2019-00077

27 Ikefuama, E. C. et al. Improved Locomotor Recovery in a Rat Model of Spinal Cord Injury by BioLuminescent-OptoGenetic (BL-OG) Stimulation with an Enhanced Luminopsin. International Journal of Molecular Sciences 23 (2022).

28 Petersen, E. D. et al. Restoring Function After Severe Spinal Cord Injury Through BioLuminescent-OptoGenetics. Frontiers in Neurology 12 (2022).

29 Iwano, S. et al. Single-cell bioluminescence imaging of deep tissue in freely moving animals. Science 359, 935–939 (2018). 10.1126/science.aaq1067

30 Inagaki, S. et al. Imaging local brain activity of multiple freely moving mice sharing the same environment. Sci Rep 9, 7460 (2019). 10.1038/s41598-019-43897-x

31 Su, Y. et al. An optimized bioluminescent substrate for non-invasive imaging in the brain. Nature Chemical Biology 19, 731–739 (2023). 10.1038/s41589-023-01265-x

32 Oh, Y. et al. An orange calcium-modulated bioluminescent indicator for non-invasive activity imaging. Nature chemical biology 15, 433–436 (2019). 10.1038/s41589-019-0256-z

33 Tian, X. et al. A luciferase prosubstrate and a red bioluminescent calcium indicator for imaging neuronal activity in mice. Nature Communications 13, 3967 (2022). 10.1038/s41467-022-31673-x

34 Lambert, G. G. et al. CaBLAM! A high-contrast bioluminescent Ca2+ indicator derived from an engineered Oplophorus gracilirostris luciferase. bioRxiv, 2023.2006.2025.546478 (2023). 10.1101/2023.06.25.546478

35 Petersen, E. D. et al. Bioluminescent Genetically Encoded Glutamate Indicators for Molecular Imaging of Neuronal Activity. ACS Synth Biol (2023). 10.1021/acssynbio.2c00687

36 Theer, P. & Denk, W. On the fundamental imaging-depth limit in two-photon microscopy. Journal of the Optical Society of America A 23, 3139–3149 (2006). 10.1364/JOSAA.23.003139

37 Raghuram, A. et al. Determining the Depth Limit of Bioluminescent Sources in Scattering Media. bioRxiv, 2020.2004.2021.044982 (2020). 10.1101/2020.04.21.044982

38 Karstens, T., Staufer, T., Buchin, R. & Grüner, F. Quantitative Assessment on Optical Properties as a Basis for Bioluminescence Imaging: An Experimental and Numerical Approach to the Transport of Optical Photons in Phantom Materials. Sensors 23, 6458 (2023).

39 Bakker, G. J. et al. Intravital deep-tumor single-beam 3-photon, 4-photon, and harmonic microscopy. Elife 11 (2022). 10.7554/eLife.63776

40 Svoboda, K. & Yasuda, R. Principles of two-photon excitation microscopy and its applications to neuroscience. Neuron 50, 823–839 (2006). 10.1016/j.neuron.2006.05.019

41 Ntombela, L., Adeleye, B. & Chetty, N. Low-cost fabrication of optical tissue phantoms for use in biomedical imaging. Heliyon 6, e03602 (2020). 10.1016/j.heliyon.2020.e03602

42 Szu, J. I. et al. Thinned-skull Cortical Window Technique for In Vivo Optical Coherence Tomography Imaging. JoVE, e50053 (2012). doi:10.3791/50053

43 Peipei, L., Zhirong, Z., Chang-Chun, Z. & Guang, N. Specific absorption spectra of hemoglobin at different PO2 levels: potential noninvasive method to detect PO2 in tissues. Journal of Biomedical Optics 17, 125002 (2012). 10.1117/1.JBO.17.12.125002

44 Spiess, E. et al. Two-photon excitation and emission spectra of the green fluorescent protein variants ECFP, EGFP and EYFP. Journal of Microscopy 217, 200–204 (2005). 10.1111/j.1365-2818.2005.01437.x

45 Kim, T. J., Türkcan, S. & Pratx, G. Modular low-light microscope for imaging cellular bioluminescence and radioluminescence. Nature Protocols 12, 1055–1076 (2017). 10.1038/nprot.2017.008

46 Werley, C. A., Chien, M.-P. & Cohen, A. E. Ultrawidefield microscope for high-speed fluorescence imaging and targeted optogenetic stimulation. Biomedical Optics Express 8, 5794–5813 (2017). 10.1364/boe.8.005794

47 Morales-Curiel, L. F. et al. Volumetric imaging of fast cellular dynamics with deep learning enhanced bioluminescence microscopy. Communications Biology 5, 1330 (2022). 10.1038/s42003-022-04292-x

